# Room-Temperature Aerosol Dehydration of Green Fluorescent Protein

**DOI:** 10.1101/2024.09.17.613563

**Authors:** Zehao Pan, Junshi Wang, Howard A. Stone, Maksim Mezhericher

## Abstract

Rapid Room-Temperature Aerosol Dehydration (RTAD) is a novel, scalable drying technology for powderization and thermal stabilization of pharmaceutical drug products. Compared to conventional spray drying, RTAD utilizes sub-20 *µ*m droplets that evaporate rapidly at room temperature, thereby reducing drying-induced stresses for thermally sensitive biologics. In this study, we employed Green Fluorescent Protein (GFP) as a model biological molecule to optimize the RTAD system design and process parameters. We investigated the effects of droplet size, multiphase flow patterns in the drying chamber, and usage of polysorbate 20 as a model surfactant on GFP fluorescence after drying and powder reconstitution. The experiments demonstrated that the presence of polysorbate 20 in the formulation significantly influenced GFP fluorescence intensity, especially for smaller droplets. The numerical studies using Computational Fluid Dynamics simulations revealed that the intensity of GFP fluorescence in the produced dry powders was dependent on the patterns of multiphase flow in the drying chamber. Non-axisymmetric flows and closed circulating streamlines near the drying gas inlet negatively impacted the GFP fluorescence intensity. Through iterative optimization of chamber design, process parameters, and feedstock formulation, we achieved the recovery (compared to the initial samples) of GFP fluorescence intensity exceeding 96% in the obtained dry powders. This work establishes GFP as a sensitive model biologic and its fluorescence intensity as a powerful tool for rapidly assessing bioactivity following dehydration. The insights gained from this study have broad implications for the design and scale-up of room-temperature drying technologies, which can potentially transform the production of dry powder biopharmaceuticals.

## 1. Introduction

Dehydration of biological pharmaceutical products is a critical step in drug product manufacturing for dry powder inhalation, intranasal drug delivery, and vaccine stabilization. Due to the sensitivity of many protein biopharmaceuticals to temperature, we leveraged our prior fundamental studies on liquid atomization and aerosol generation (*1, 2*) and developed a method for laboratory-scale Rapid Room-Temperature Aerosol Dehydration (RTAD) (*3, 4*). Compared to traditional spray drying, RTAD leverages much smaller droplet sizes (sub-20 *µ*m) to enhance the rate of solvent evaporation rather than relying on the elevated temperature of a drying gas. In our recent study, we demonstrated RTAD offers a system for room-temperature dehydration of liquid biopharmaceutical formulations of antigen-binding fragment (Fab) of monoclonal antibody with reduced product degradation compared to traditional spray drying (*4*).

Despite the reduction of thermal stresses with the help of RTAD’s room-temperature process, biological products could still be subject to interfacial and shear stresses during the atomization and drying steps in RTAD (*5*). In this work, we used Green Fluorescent Protein (GFP) as a model biological molecule to assess the efficacy of our approach for retaining biological activity and to optimize the RTAD process and feedstock formulation, including the conditions of the drying gas. If GFP experiences a structural variation during the process, such as unfolding or aggregation, the excitation and emission spectrum of GFP would be altered, resulting in shifts in fluorescence intensity at a fixed wavelength of the measurement (*6–8*). Thus, by comparing the fluorescence intensity of GFP at fixed wavelength before and after dehydration, we can assess the effect of the external stresses induced by drying of GFP.

Through this approach, we identified three major conditions in RTAD that can affect the GFP fluorescence intensity, including the use of surfactant, droplet size, and the recirculation patterns of the drying gas in the RTAD chamber. Our work demonstrates that GFP can be used as a model biological molecule to test the design of the feedstock formulation, atomization into droplets, and patterns of multiphase flow in the RTAD process and, in general, other spray drying systems.

## 2. Materials and Methods

### 2.1. Materials

Mannitol and polysorbate 20 were purchased from Sigma Aldrich (USA). 1X Tris-EDTA buffer (pH 7.4), which contains 10 mM tris(hydroxymethyl)aminomethane and 1 mM ethylenediaminetetraacetic (EDTA) acid was purchased from Fischer Scientific (USA). Technical grade GFP was purchased from Protein Mods (Wisconsin, USA). The PVDF filters were purchased from Tisch Scientific (Ohio, USA). In all feedstock formulations, 1X Tris-EDTA buffer, 7.5 wt% mannitol, and 0.03 mg/ml GFP were used. When surfactant was present, 0.4 mg/ml polysorbate 20 was added to the feed-stock.

### 2.2. Measurement of droplet size distribution

The Rapid Room-Temperature Aerosol Dehydration system utilized a proprietary two-fluid nozzle (INAEDIS, INC., Princeton, NJ, USA) (*9*) capable of generating fine droplet sprays with diameters typically ranging from 0.1 to 20 *µ*m. This droplet size range is characteristic of aerosols rather than conventional spray nozzles, enabling more efficient dehydration at room temperature. The droplet size distribution of the atomizer was measured with laser diffraction (Spraytec, Malvern Panalytical, UK) using the previously described procedure (*2*). The nominal measurement accuracy of the Malvern Spraytec device is better than ± 1% for the median diameter of a droplet size distribution. For each experimental run to measure the droplet size distribution, aerosol was generated for a minimum of one minute under quasi-steady state conditions, ensuring consistent and reliable data collection. This approach allowed for the establishment of a stable aerosol output and provided sufficient time for accurate sampling and analysis. The laser beam crossed the aerosol flow at a right angle and the samplings of the droplet size distribution were performed 6 cm from the atomizer. The scattered signal of the laser beam was recorded with a frequency of 1 Hz, allowing for averaging 20-30 droplet size distribution profiles for each operating condition.

Deionized water (Milli-Q, 18.2 MΩ · cm) served as the test liquid in our experimental investigation of the volume-weighted median droplet diameter (Dv50) and Sauter mean diameter (D_32_) of the generated droplet aerosols. We systematically examined how these key droplet size parameters varied in response to changes in the atomizing gas pressure and liquid feed flow rate supplied to the atomizing nozzle, providing insights into the atomization process dynamics.

### 2.3. Measurement of particle morphology and size distribution

Particle morphology and size distribution were analyzed using scanning electron microscopy (SEM, Quanta 200 FEG Environmental-SEM). To preserve sample integrity, powders collected on the filter were carefully applied to SEM stubs within a humidity-controlled glove box. Prior to imaging, the samples underwent iridium coating via plasma sputtering (Leica EM ACE600 Sputter Coater) to enhance conductivity and image quality. SEM imaging was conducted in low-vacuum mode to minimize sample distortion. For statistical robustness, the particle size distribution analysis incorporated measurements from over 500 individual particles, ensuring a comprehensive representation of the sample population.

### 2.4. Measurement of powder residual moisture content

Residual moisture content of the powder was determined using a thermogravimetric analyzer (PerkinElmer TGA-8000). To ensure consistent initial conditions, the powder was stored overnight in a humidity-controlled chamber maintained at room temperature and less than 5% relative humidity. Subsequently, a 5 to 10 mg sample was weighed precisely and placed in a ceramic sample pan (PerkinElmer). The analysis protocol involved loading the sample into the furnace, heating it to 110 °C at a rate of 20 °C/min, and then maintaining this temperature for two hours. The weight loss attributed to moisture evaporation was continuously monitored and ultimately expressed as a percentage of the initial sample weight.

### 2.5. Characterization of GFP fluorescence

To assess the preservation of GFP functionality after drying, we compared the fluorescence intensity of the reconstituted powder to that of the original feedstock solution. The dried powder was weighed precisely and dissolved in deionized water to match the feedstock concentration of 0.03 mg/ml GFP used in the RTAD process. Both the feedstock and reconstituted powder solutions were pipetted into five wells of a half-area 96-well plate (Corning, NY, USA), with each well containing 100 *µ*l. Fluorescence intensities of the feedstock (*I*_f_) and reconstituted powder (*I*_p_) were measured using a SpectraMax i3x plate reader (Molecular Devices, San Jose, CA). The fluorescence intensity ratio (*I*_p_*/I*_f_) was then calculated to quantify the retention of GFP activity. To account for experimental variability, the standard deviation of this ratio was determined using the statistical method for the ratio of two random variables, as described in reference (*10*):

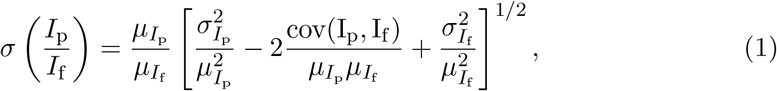

where 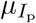 and 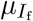 are the means, 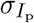 and 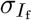 are the standard deviations, and cov(*I*_p_, *I*_f_) is the covariance of *I*_p_ and *I*_f_.

### 2.6. Room-temperature aerosol dehydration setup, drying chamber design and operating parameters

The functional principle and schematic diagram of Rapid Room-Temperature Aerosol Dehydration process are described in detail in our previous papers (*3, 4*). In this study, we used the same RTAD setup as in (*4*), and investigated the design of drying chamber. For this purpose, as RTAD chamber we utilized a stainless steel cylinder with inner diameter of 247 mm and length of 610 mm. A 0.45 *µm* polyvinylidene fluoride (PVDF) filter placed on a filter plate at the chamber’s bottom collected the produced dried particles. The top cover of the chamber housed both the drying gas inlet and atomizing nozzle. Nitrogen served as the drying gas, supplied at a fixed rate of 50 SLPM (L/min at standard conditions: 100 kPa, 0 °C) controlled by an Alicat Scientific controller (AZ, USA). An ELVEFLOW microfluidic pump (Paris, France) regulated the atomizing gas pressure, with flow rates ranging from 5 to 14 SLPM depending on atomization pressure. The Reynolds numbers for the drying and atomizing gases were 9000 and 10500, respectively, calculated as Re = *uD/ν*, where *u* was gas velocity at inlet or nozzle, *D* was the inlet or nozzle diameter, and *ν* was the kinematic viscosity of the gas (nitrogen) at 22 °C. The liquid feedstock flow rate was maintained at 0.5 ml/min, limited by the drying gas system’s capacity to ensure complete product drying (detailed discussion is provided in Section 3.2). Each RTAD experiment utilized a 10 ml solution batch, with 8 ml used for dehydration and 2 ml retained as an initial reference.

Three distinct drying chamber configurations, each featuring a unique drying gas inlet design, were constructed and investigated, as illustrated in Figure 1. Configuration A employed a drying gas supply port positioned 100 mm from the top plate’s center, delivering gas at an inlet velocity of 6.6 m/s. In configuration B, the drying gas was centrally introduced into a cylindrical plenum chamber. Configuration C, a modification of B, incorporated a plenum diffuser with an 8° total divergence angle (*11*). Both configurations B and C operated with an inlet gas velocity of 24.9 m/s. These design variations allowed for an analysis of gas flow dynamics and their impact on the RTAD process efficiency.

**Figure 1.**
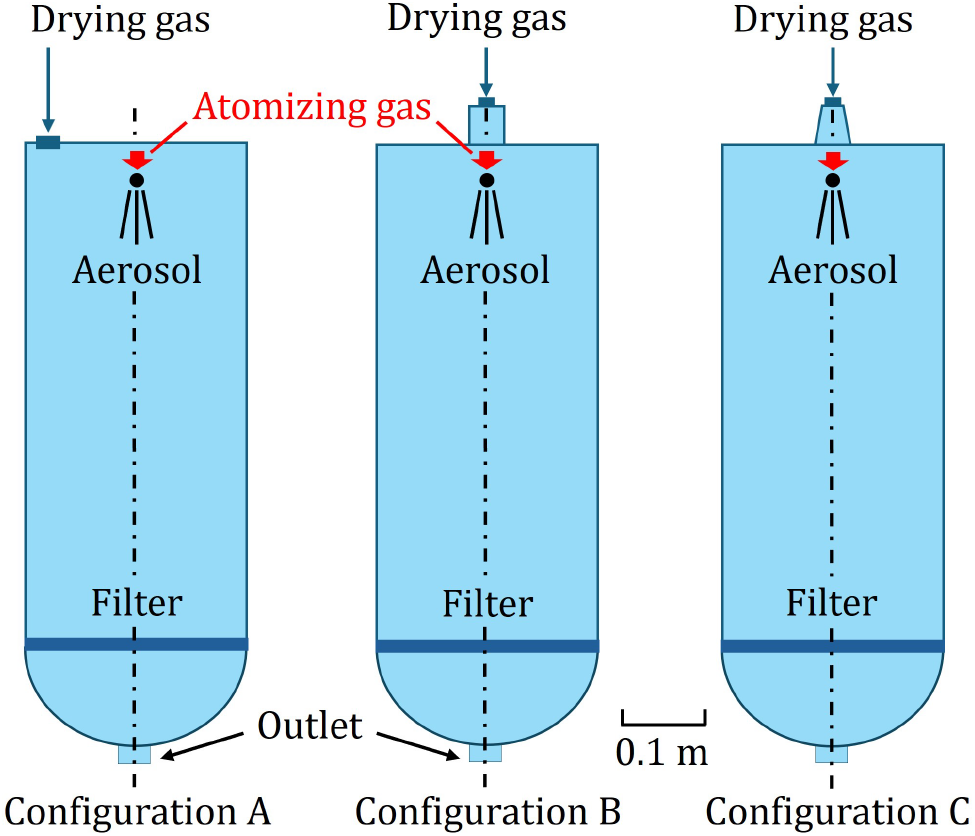
Schematic of drying chamber configurations used in the study of the laboratory-scale RTAD process. In configuration A, the drying gas was injected from a port placed 100 mm away from the center of the top plate, where the drying gas inlet velocity was 6.6 m/s. In configuration B, the drying gas was introduced from the center into a cylindrical plenum chamber. Compared to configuration B, configuration C has a diffuser in the plenum with a total divergence angle of 8° (*11*). All three chambers have an atomization nozzle placed in the center with flow of an atomizing gas to generate droplets. Dried particles are collected at the filter.

Table 1 summarizes the experimental conditions investigated in this study. Droplet size variations were achieved by manipulating two key parameters: the atomizing gas pressure and the solution surface tension through the addition of polysorbate 20. To evaluate the impact of the drying gas inlet design, the above-mentioned three drying chamber configurations were studied using identical feedstock formulations and droplet sizes in experimental runs 4, 5, and 6. This systematic approach allowed for an assessment of the influence of both droplet characteristics and drying chamber geometry on the RTAD process performance.

**Table 1.**
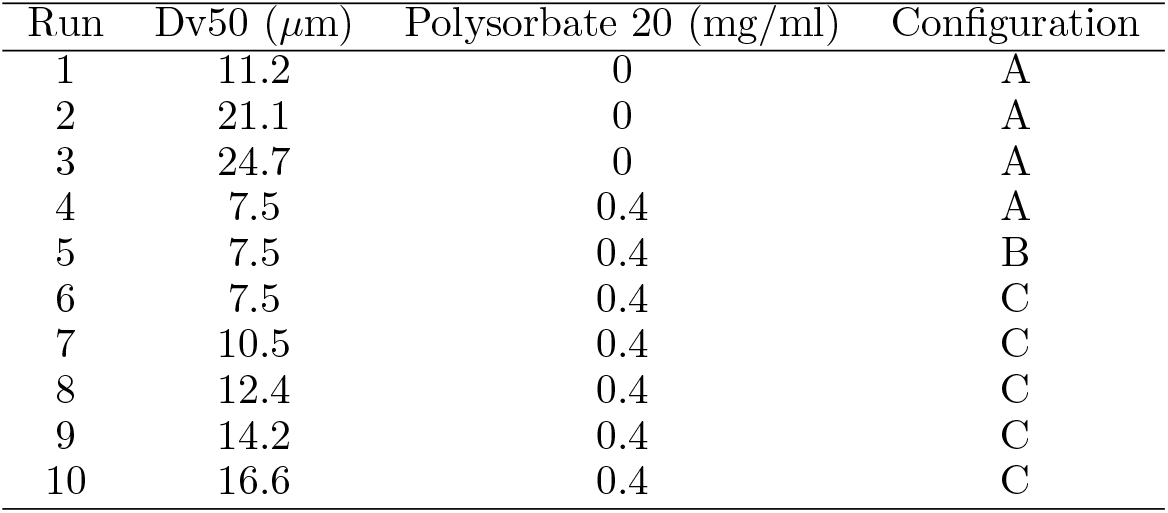
Experimental conditions for all RTAD runs. The values of Dv50 were derived from atomizing gas pressure using the experimental relationship shown in Figure 3 and adjusted for the presence of polysorbate 20 by applying equation (2). Feedstock composition: 0.03 mg/ml GFP, 75 mg/ml mannitol, 1X Tris-EDTA buffer.

### 2.7. Multiphase flow modeling using Computational Fluid Dynamics

We conducted full-scale, three-dimensional, steady-state Computational Fluid Dynamics (CFD) modeling and numerical simulations of the RTAD processes for three drying chamber configurations (Figure 1): Configuration A (side port inlet), Configuration B (center port inlet without diffuser), and Configuration C (center port inlet with diffuser). The numerical modeling and simulations were performed using ANSYS Fluent 2024 R1 software package, building upon the CFD model of the spray drying process previously developed by Mezhericher et al. (*12, 13*).

An Eulerian-Lagrangian model was employed to simulate the multiphase flow, treating the gas flow as a continuous phase (Eulerian approach) and droplets/particles as the discrete phase (Lagrangian approach). The gas phase was governed by the Reynolds-Averaged Navier-Stokes (RANS) and the energy conservation equations, incorporating gravity and ideal gas behavior. Viscous modeling utilized the *k*-*ϵ* turbulence model with standard wall functions. Species transport considered a nitrogen-water vapor mixture as an ideal gas. The spray of evaporating droplets was modeled using the Discrete Phase Model (DPM) by incorporating multi-component droplet injections at the nozzle outlet (Figure 2a). The experimental hollow spray cone was modeled by 500 DPM droplet injections, with size distribution simulated using the Rosin-Rammler equation (*14*). Each droplet injection comprised 25 different droplet diameters (0.5-30 *µ*m) introduced into the chamber at 22 °C, matching the droplet size distribution produced by the RTAD nozzle as measured by laser diffraction at 2 bar of atomizing gas pressure and 1 ml/min feedstock flow rate (Figure 3). As droplets traversed the RTAD chamber, carried by the mixed streams of the drying and atomizing gases, they were considered to be either “escaped” when hitting the particle filter, or “trapped” when deposited on the chamber walls.

**Figure 2.**
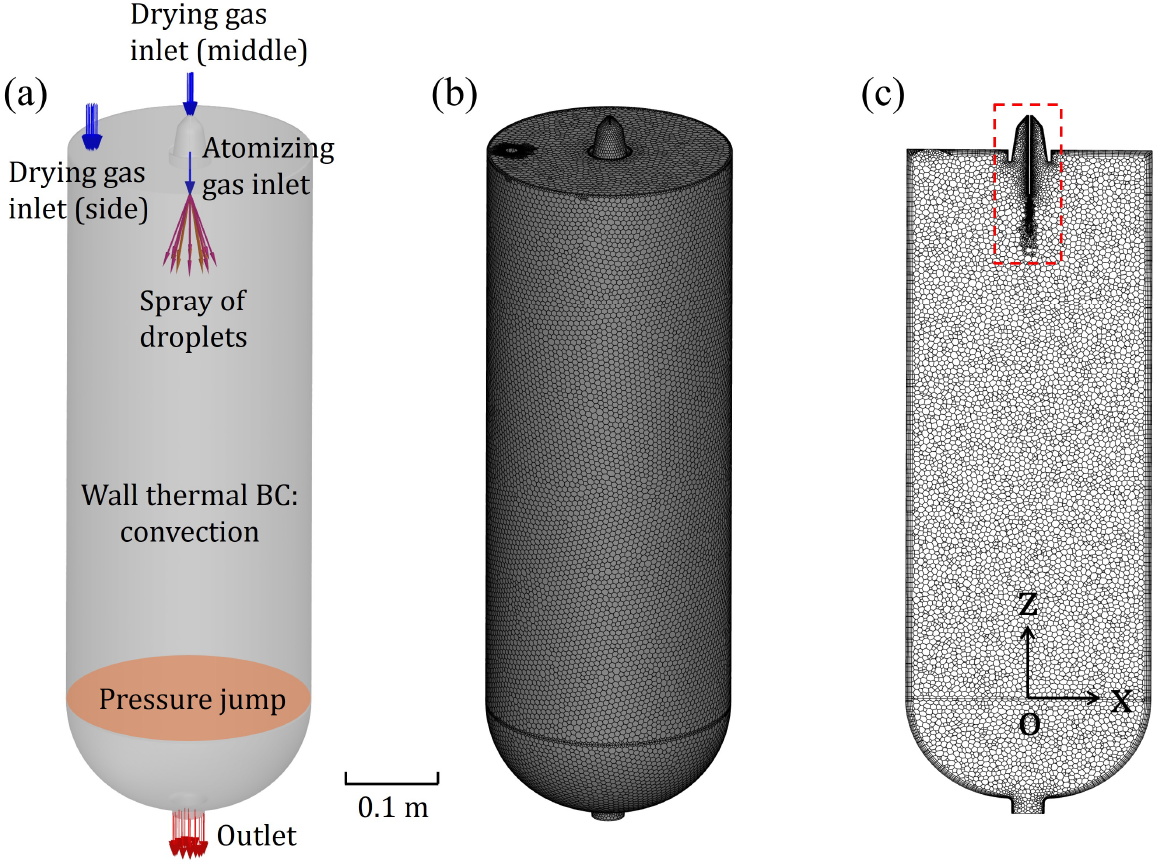
Computational modeling of the RTAD process in a laboratory-scale drying chamber. (a) Drying chamber geometry with boundary conditions (BC) including drying gas inlets at the center and side, atomizing gas inlet and cone-shape droplet injections from the spray nozzle, pressure jump across the particle filter, and gas outlet of the chamber. (b) Polyhedral numerical mesh on the internal chamber surface. (c) Polyhedral numerical mesh on the middle vertical cross section plane of the computational domain. The numerical mesh was refined by using the Adaptive mesh refinement (AMR) algorithm, targeting mesh cells where the gradient of the magnitude of gas velocity was higher than 15% of the global maximum of the gas velocity gradient.

**Figure 3.**
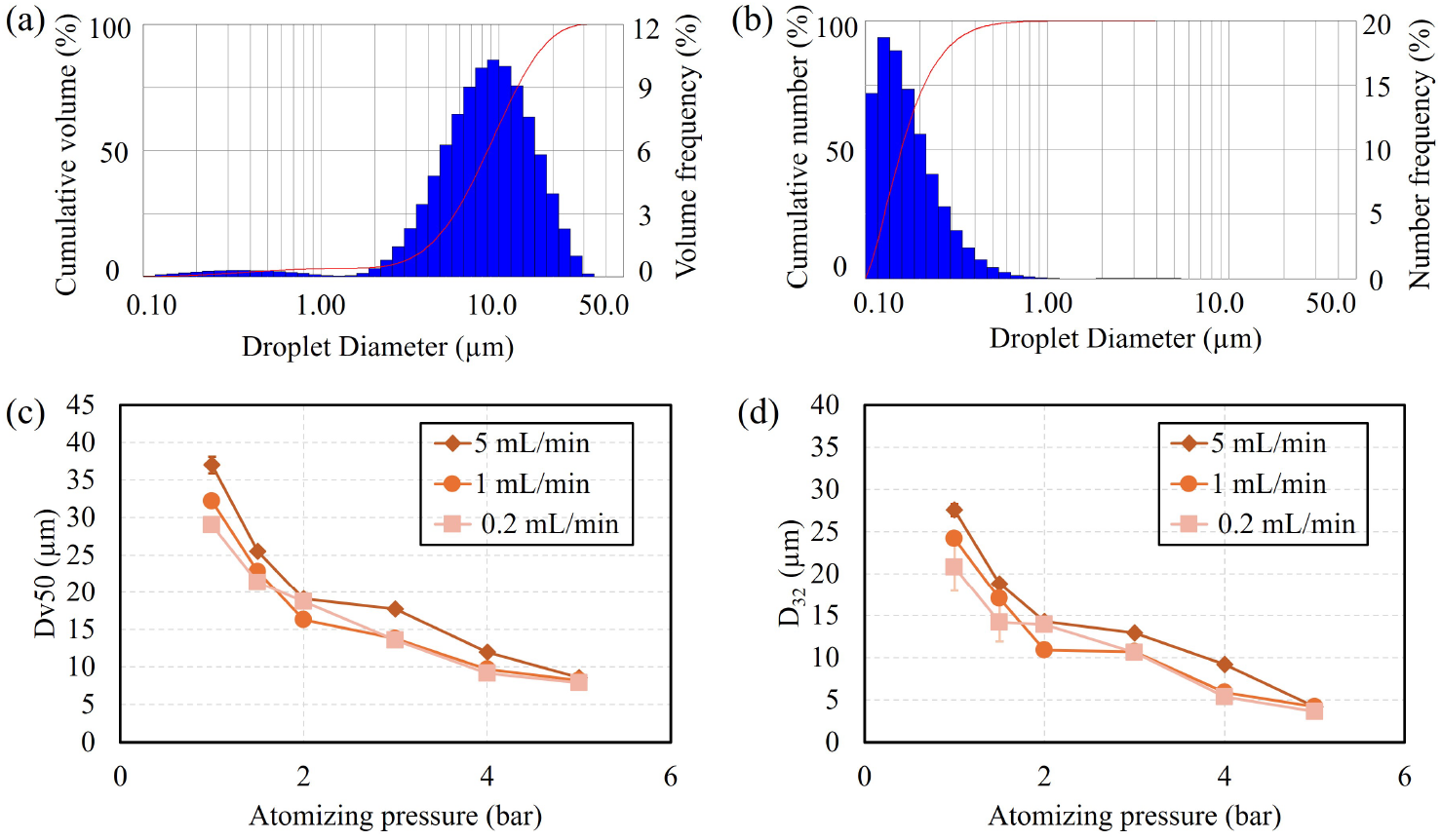
Characteristic diameters of water droplets generated using our proprietary liquid atomization technique (*9*) and analyzed via laser diffraction. (a) Volume-weighted and (b) number-weighted size distributions of aerosol droplets produced at 5 ml/min water flow rate and 5 bar atomization pressure. (c) Dv50 (volume median diameter) and (d) D_32_ (Sauter mean diameter) of atomized droplets as functions of atomizing gas pressure at various liquid feed flow rates. Error bars indicate standard deviations from 10 replicate measurements.

The boundary conditions included mass flow inlets for both drying and atomizing gases (Figure 2a). The drying gas was introduced via a side port for configuration A and a center port for configurations B and C (Figure 1). Atomizing gas was injected at the nozzle exit in all configurations. Both atomizing and drying gases were set as pure nitrogen at 22 °C, with flow rates of 50 SLPM (drying) and 10 SLPM (atomizing). A pressure jump boundary condition was applied at the particle collection filter, with a pressure outlet at the chamber exit (Figure 2a). The chamber wall was modeled as a no-slip thermally conducting surface with convective thermal conditions, assuming 22 °C ambient temperature.

Droplet drying kinetics were simulated using a two-component mixture model: evaporating water and non-evaporating solid material. For simplicity, both components were assigned water’s density, thermal conductivity, and heat capacity. Droplets with water content below 5 wt% were treated as non-evaporating spherical particles in subsequent calculations.

The computational domain utilized a polyhedral mesh (Figure 2b) with adaptive mesh refinement (AMR) in regions of high velocity gradients (Figure 2c). Mesh size studies ensured numerical stability, solution accuracy, and mesh independence. Detailed computational setup and mesh-independence studies are provided in Appendices A and B.

## 3. Results and Discussion

### 3.1. Droplet size distribution

The measurements shown in Figures 3(a) and (b) illustrate typical volume-weighted and number-weighted size distributions of spray droplets generated by the proprietary RTAD atomizing nozzle as measured by laser diffraction. The volume median diameter, Dv50, represents the diameter below which 50% of the total aerosol or spray volume is contained (*15*). The Sauter mean diameter, D_32_, reflects the volume-to-surface-area ratio of the spray or aerosol equivalent to a sphere with diameter D_32_ (*15*). The results reported in Figures 3 (c) and (d) depict Dv50 and D_32_ as functions of atomizing pressure at three distinct water flow rates.

The droplet sizes exhibit minimal dependence on liquid feed flow rate, demonstrating the atomizer’s scale-up potential. A 25-fold increase in liquid flow rate (0.2 mL/min to 25 mL/min) results in less than 20 % increase in Dv50 at 2 bar atomization pressure, and less than 10 % at 5 bar.

In pharmaceutical formulations such as Avastin (*16*), 0.4 mg/ml of polysorbate 20 is typically included into the aqueous solutions. This addition reduces the surface tension of solution to 37 mN/m (*17*), approximately half that of water. For low-viscosity solutions in atomization regimes where viscous forces are negligible compared to inertial and surface tension forces (Ohnesorge number ≪ 1), the droplet diameter is proportional to surface tension (*18*). The Ohnesorge number is defined as 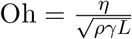, where *η* is liquid viscosity, *ρ* is liquid density, *γ* is surface tension, and *L* is the typical length scale (e.g., the droplet diameter). This relationship enables the calculation of the volume-weighted median drop diameter for polysorbate 20 solutions based on the water Dv50 curve, using the equation:

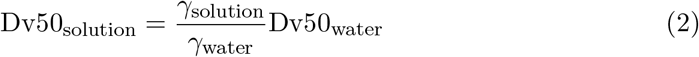

### 3.2. Single droplet drying kinetics and drying capacity

In this study, we employed our previously developed analytical model of single droplet drying kinetics (*12, 13, 19*) to calculate and compare evaporation times for droplets of varying diameters. The calculation uses equation 20 from (*13*).

For the experiments outlined in Table 1, droplet evaporation times were computed under conditions of 5% relative humidity (RH) in the surrounding gas and a relative gas-droplet velocity of 1 m/s (Figure 4a). The Dv50 values from each experiment (Table 1) are plotted as discrete points. Utilizing the same single droplet drying kinetics model (*12, 13, 19*), we calculated the equilibrium evaporation temperature of droplets during the drying process. Figure 4 illustrates the strong dependence of droplet temperature during dehydration on the surrounding gas’s relative humidity, comparing scenarios with 5% and 30% RH. The latter RH range is characteristic of the RTAD processes summarized in Table 1. Notably, the droplet evaporation temperature during drying closely approximates the wet bulb temperature and remains independent of droplet diameter (*19*). This analysis provides crucial insights into the drying dynamics and thermal conditions experienced by droplets in the RTAD process, facilitating optimization of process parameters for efficient dehydration while minimizing thermal stress on sensitive biological materials.

**Figure 4.**
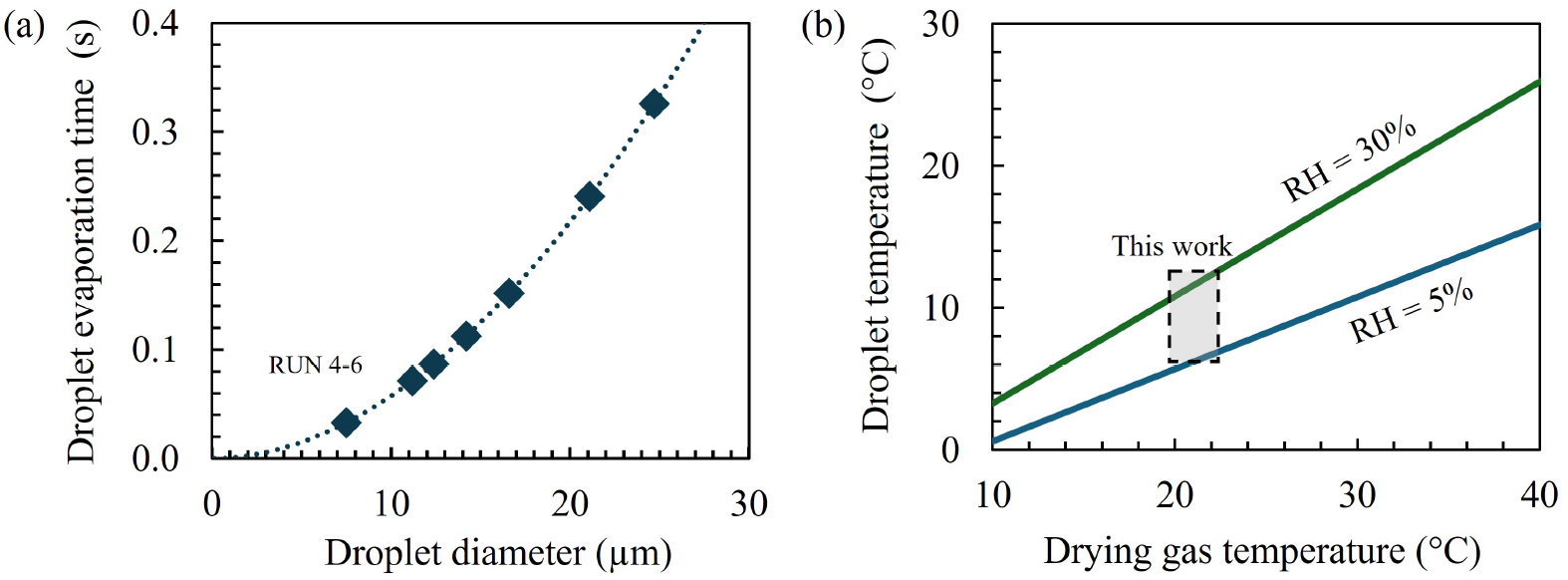
Single droplet drying kinetics for RTAD experiments summarized in Table 1. (a) Droplet evaporation time as a function of droplet size. Dotted line represents calculations using equation 20 from (*13*), assuming 5% relative humidity (RH), 22 °C gas temperature and 1 m/s relative gas-droplet velocity. Points indicate calculated data for specific Dv50 values from Table 1. (b) Droplet temperature versus drying gas temperature at various values of gas relative humidity. Lines show theoretical curves based on wet bulb temperature at fixed RH values. Shaded square denotes the operating regime of the RTAD system in this study.

The drying capacity of RTAD process, i.e., the mass flow rate of evaporating solvent from the feedstock 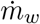, is equal to the mass flow rate of vaporized solvent at the outlet of the drying chamber 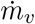, i.e. 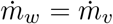. The drying capacity is limited by the drying gas flow rate 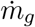 and relative humidity *φ* of gas-vapor mixture at the outlet of RTAD chamber. According to Dalton’s law, the measurable total pressure *p*_*tot*_ at the outlet consists of *p*_*v*_, the partial pressure of water vapor, and *p*_*g*_, the partial pressure of drying gas:

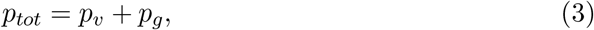

where *p*_*v*_ is a function of *φ* and the saturation vapor pressure *p*_*sat*_ at gas temperature *T*_*g*_:

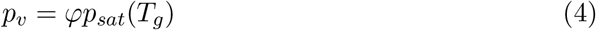

When RTAD operates in a steady state, for a gas-vapor mixture at the outlet of the drying chamber we can assume that 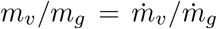, where *m*_*v*_ and *m*_*g*_ are, respectively, masses of water vapor and drying gas. Using the ideal gas law for both vapor and drying gas at the outlet of RTAD chamber, we can obtain:

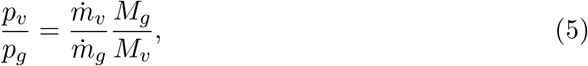

where *M*_*v*_ and *M*_*g*_ are, respectively, the molar mass of water and nitrogen, i.e., 18.015 g/mol and 28.013 g/mol. Combining equations (3-5) and bearing in mind that 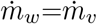, the relationship between mass flow rate of evaporating solvent in feedstock 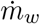 and mass flow rate of drying gas 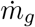 is given by:

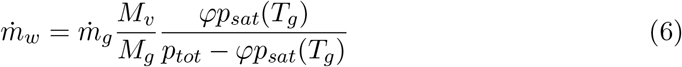

In this study, to keep the collected product on the particle filter dry, we targeted the relative humidity of the gas-vapor mixture at the chamber outlet to be less than *φ* = 0.25. Based on the equation (6), we selected 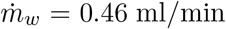, so that the total supplied liquid feedstock flow rate was maintained at 0.5 ml/min.

### 3.3. Particle morphology and size distribution

Analysis of powder particles produced across all experiments consistently revealed smooth spherical morphology of the particles. A representative SEM micrograph of the powder produced in the experimental run 7 is shown in Figure 5a. The rapid drying kinetics of sub-20 *µ*m droplets containing dissolved mannitol promote supersaturation and high nucleation rates, resulting in crystal formations too small to be discerned at the given SEM magnification (*20*). Notably, some particles exhibit visible holes, indicating potential hollow internal structures. Figure 5b depicts the number-weighted particle size distribution, derived from particle counts in Figure 5a, with a count median diameter of 1.5 *µ*m.

**Figure 5.**
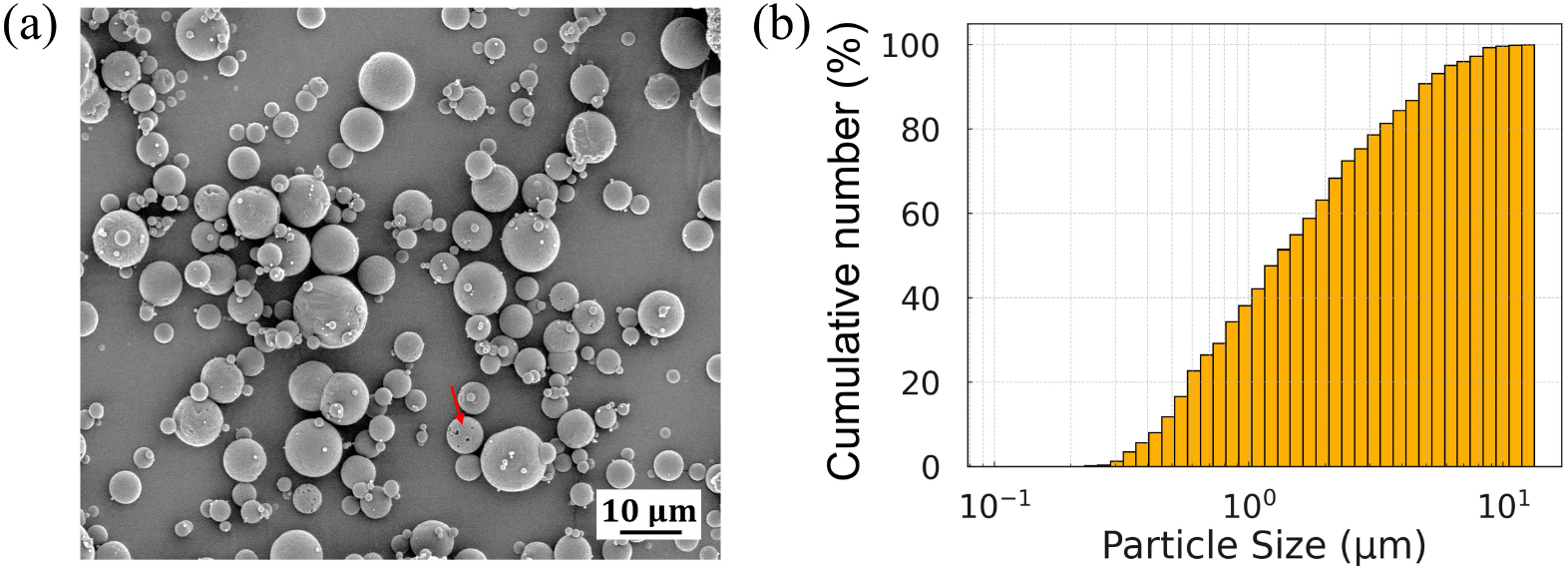
(a) SEM micrograph of particles produced in run 7. Red arrow points to holes in a particle. (b) Cumulative number-weighted size distribution of particles depicted in panel (a). The distribution curve illustrates the percentage of particles smaller than a given diameter.

The observed morphology of the obtained particles can be explained by analyzing a droplet as a diffusion-advection system with receding boundary. For typical 10 *µ*m droplet, after introducing into the drying chamber the relaxation time and time to reach terminal velocity are respectively 0.3 ms and 0.9 ms (*15*). Those time scales are much shorter compared to evaporation time of around 65 ms calculated for this droplet diameter by using the theoretical model of Mezhericher et al. (*19*). Therefore, neglecting shear-induced internal circulation in droplet after it reaches the relaxation, we consider a competition between evaporation-induced shrinkage of droplet radius and diffusion of solids in droplet as two phenomena determining the morphology of formed particles. The ratio between the rates of these two processes is described by the Peclet number, *Pe* (*21, 22*):

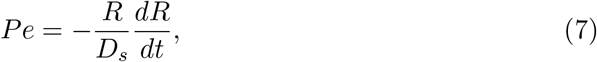

where *R* is droplet radius and *D*_*s*_ is the diffusivity of solids in droplet solution. Using the theoretical model of single droplet drying by Mezhericher et al. (*19*) to determine 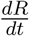, we can obtain:

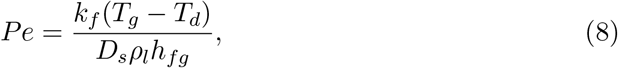

where *T*_*g*_ and *T*_*d*_ are, respectively, the drying gas and droplet temperatures, *k*_*f*_ is the thermal conductivity of fluid around the droplet surface, and *ρ*_*l*_ and *h*_*fg*_ are, respectively, the density and latent heat of vaporization of the solvent. Using for water and water vapor *ρ*_*l*_ = 1000 kg/m^3^, *k*_*f*_ = 0.025 W/(m·K), *h*_*fg*_ = 2478 kJ/kg, *D*_*s*_ = 6.7×10^−10^ m^2^/s for mannitol (*23*) and *D*_*s*_ = 9.0×10^−11^ m^2^/s for GFP (*24*), and *T*_*g*_ = 22 °C and *T*_*d*_ = 11 °C, we obtain Peclet numbers for mannitol and GFP as 0.15 and 1.1, respectively. This indicates that diffusion of GFP and evaporation-induced shrinkage of droplet occur over equal times, while the diffusive transport of mannitol dominates over both phenomena. We conclude that molecules of mannitol have time to redistribute by diffusion throughout the evaporating droplet, thus yielding relatively dense spherical dried particles (see Figure 5a), consisting of mannitol matrix incorporating GFP.

Additionally, from equation (8) we find that *Pe* ∝ (*T*_*g*_ − *T*_*d*_) and *Pe* does not directly depend on droplet diameter. Therefore, we can expect that for the same feedstock formulation, Peclet number of spray drying will be larger by a factor 10 of that of RTAD, because of the 10x greater (*T*_*g*_ − *T*_*d*_) temperature differences in spray drying (∼100 °C or more) versus RTAD (∼10 °C). The 10x smaller value of *Pe* for RTAD process explains the considerable difference between the morphologies of spray dried particles (crumpled non-spherical) and particles of RTAD (spherically shaped) observed in our previous study (*4*).

### 3.4. Residual moisture content of produced powders

For all experiments, the residual moisture content of the produced powders, as measured by the loss of weight method, consistently ranged from 1.5% to 1.9%. This narrow range demonstrates the capability of the RTAD process to achieve uniform and low moisture content across various experimental conditions, which is a critical factor for ensuring product stability and quality.

### 3.5. GFP fluorescence spectra before and after dehydration

To assess the biological activity of GFP, we measured and compared the excitation and emission spectra of the reconstituted powder against those of the unprocessed feedstock solution. Figure 6a presents representative data from this analysis. Notably, both the reconstituted powder and feedstock solutions exhibited identical spectral characteristics: an absorption peak at 475 nm and an emission peak at 505 nm. This consistency in spectral profiles indicates that the RTAD process did not induce any significant structural changes in the GFP molecule that would manifest as spectral shifts. The preservation of these key spectral features suggests that the GFP retained its native conformation and fluorescent properties throughout the dehydration and reconstitution processes.

**Figure 6.**
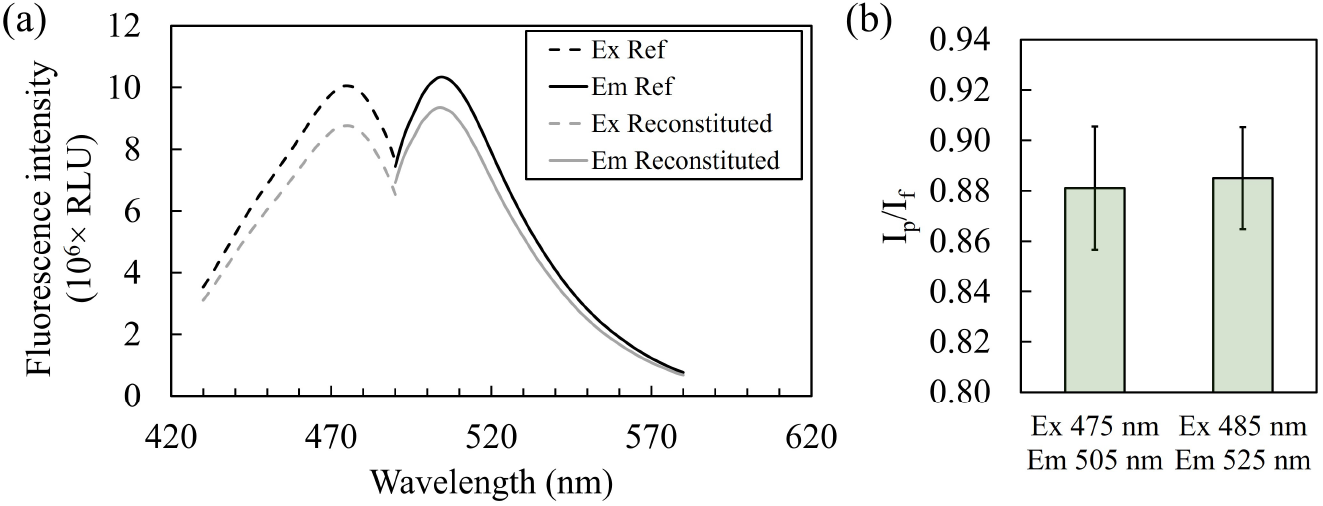
(a) Excitation and emission spectra of GFP before and after drying from experimental run 4 measured in Relative Light Units (RLU). Dashed lines represent excitation spectra, and solid lines represent emission spectra. Excitation spectra were obtained by scanning excitation wavelengths with a fixed emission wavelength at 525 nm. Emission spectra were obtained by scanning emission wavelengths with a fixed excitation wavelength at 450 nm. The intensity standard deviation is within ±6% of the intensity magnitude. (b) Ratio of GFP fluorescence intensity (*I*_p_*/I*_f_) after drying (*I*_p_) to before drying (*I*_f_), measured at Ex 475 nm - Em 505 nm and Ex 485 nm - Em 525 nm. This comparison highlights the preservation of GFP’s fluorescent properties through the drying process.

Two pairs of excitation (Ex) and emission (Em) wavelengths were employed for analysis: 1) 475 nm/505 nm and 2) 485 nm/525 nm. The latter pair (485 nm/525 nm) corresponds to the excitation wavelength used in commercial fluorescence intensity detection cartridges for the SpectraMax i3x system. Figure 6b illustrates the results for both wavelength pairs. Notably, both options yielded equivalent GFP fluorescence intensity ratios (*I*_p_*/I*_f_), where *I*_p_ represents the intensity after drying and *I*_f_ before drying. Consequently, these two Ex/Em wavelength pairs (475 nm/505 nm and 485 nm/525 nm) were used to quantify the variation in fluorescence intensity after drying across all RTAD runs, ensuring consistency and comparability in the assessment of the preservation of GFP activity.

### 3.6. Ratio of GFP fluorescence intensity I_p_/I_f_ as a function of the inlet configuration of drying gas

The “drying gas inlet” panels in Figure 7 illustrate the gas flow patterns across the drying chamber and near the drying gas inlet for each configuration. In configuration A, the spray of aerosol droplets is released outside the central drying gas flow as it enters the chamber, contrasting with configurations B and C. Configuration B exhibits strong recirculation zones at the corners of the cylindrical plenum housing the drying gas inlet. Configuration C eliminates these recirculation zones by replacing the cylindrical plenum with a conical diffuser, featuring an 8° divergence angle (*11*).

**Figure 7.**
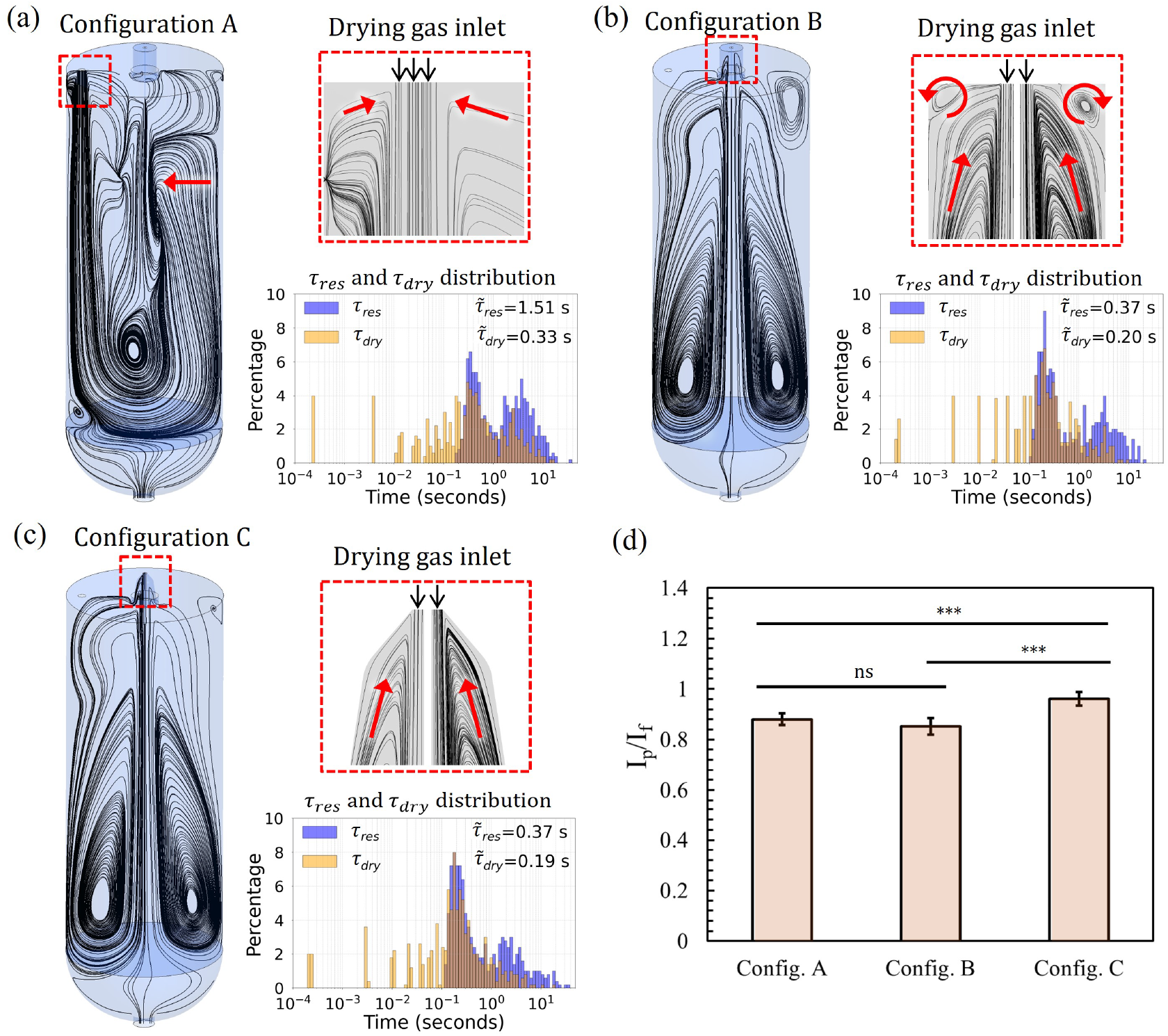
(a-c) CFD simulation of the RTAD process for drying chamber configuration A (a), B (b) and C (c). For each configuration, the panels display gas streamlines for the entire chamber, along with an enlarged view near the drying gas inlet, highlighted by a red square. Additionally, the residence time *τ*_*res*_ and drying time *τ*_*dry*_ distributions for each simulation are plotted. (d) Experimental results of *I*_p_/*I*_f_ of GFP processed with chamber configurations A, B and C. ns: not significant; ∗ ∗ ∗ : *p <* 0.001 (Student’s t-test assuming same *I*_p_/*I*_f_ between two configurations).

These distinct flow patterns significantly influence the droplet trajectories and drying conditions within the chamber, potentially affecting the final product characteristics. The drying process of a single droplet during RTAD is characterized by two key time parameters: the particle residence time (*τ*_*res*_), representing the duration a drying droplet or dried particle remains airborne, and the droplet drying time (*τ*_*dry*_), defined as the period during which a droplet retains water content above 5 wt%, the final residual water content set for this study. The “*τ*_*res*_ and *τ*_*dry*_ distribution” panels in Figures 7a-c illustrate the distributions of these time parameters. Under uniform drying conditions, *τ*_*dry*_ correlates monotonically with droplet diameter, as shown by the single droplet drying kinetics model (Figure 4a). Given the unimodal droplet size distribution (Rosin-Rammler) assumed for the RTAD atomizer, uniform drying conditions should yield a unimodal *τ*_*dry*_ distribution. Configuration C exhibits a near-unimodal *τ*_*dry*_ distribution, indicating more uniform drying conditions compared to the other configurations. Configuration A’s longer median 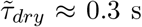, versus 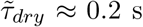 for configurations B and C, reflects its off-center drying gas inlet and centerline-located liquid atomizer, promoting larger recirculation zones and extended drying times.

As the results in Figures 7a-c reveal, the numerical modeling using CFD simulations predicted median 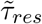 values of approximately 1.5 s for configuration A and 0.4 s for configurations B and C. The extended airborne time in configuration A suggests that dried particles may re-enter multiple times the top region close to the atomizer, experiencing repeated exposure to high gradients of velocity, humidity, temperature, pressure, and species. This also increases the likelihood of inter-particle and particle-wall collisions before particles collection by the filter.

The measures shown in Figure 7d demonstrate the ratios of GFP fluorescence intensity *I*_p_/*I*_f_ measured from the runs 4, 5, and 6. It is worth noting that the differences between the configuration C and the configurations A and B are statistically significant (p-values *<* 0.001 from Student’s t-test). The observed variations in *I*_p_/*I*_f_ can be attributed to the distinct gas flow patterns in different drying chamber configurations, as this is the only difference among the different runs. Specifically, significantly longer median *τ*_*res*_ and recirculating gas streamlines for configuration A suggest that particles may repeatedly encounter high gradients and increased inter-particle collision and wall deposition probabilities (*14*). Configuration B, compared to C, exhibits less uniform drying conditions, evident from its multimodal *τ*_*dry*_ distribution (Figure 7b and c). Additionally, configuration B features a small region of closed recirculating streamlines near the drying gas entrance, potentially trapping particles and increasing collision chances. Post-experiment observations of run 5 revealed significant particle accumulation in the cylindrical plenum region, aligning with CFD predictions of a closed recirculation zone. This recirculation may create a comminution zone, enhancing particle size reduction and surface modification through increased collision energy (*25, 26*).

### 3.7. Ratio of GFP fluorescence intensity I_p_/I_f_ as a function of droplet size and the presence of surfactant

We systematically investigated the effect of the size of atomized droplets and presence of surfactant on the fluorescence intensity of reconstituted GFP powders. In the absence of polysorbate 20, droplet size significantly influenced the fluorescence intensity of reconstituted GFP. For experimental runs 1-3, the *I*_p_/*I*_f_ ratio increased from 0.58 to 0.86 as the volume median diameter of atomized droplets increased from Dv50 = 11.2 *µ*m to 24.7 *µ*m (Figure 8). Notably, the Sauter mean diameter increased from 6.9 *µ*m to 18.4 *µ*m, indicating a 2.7-fold increase in volume-to-surface area ratio of aerosolized solution.

**Figure 8.**
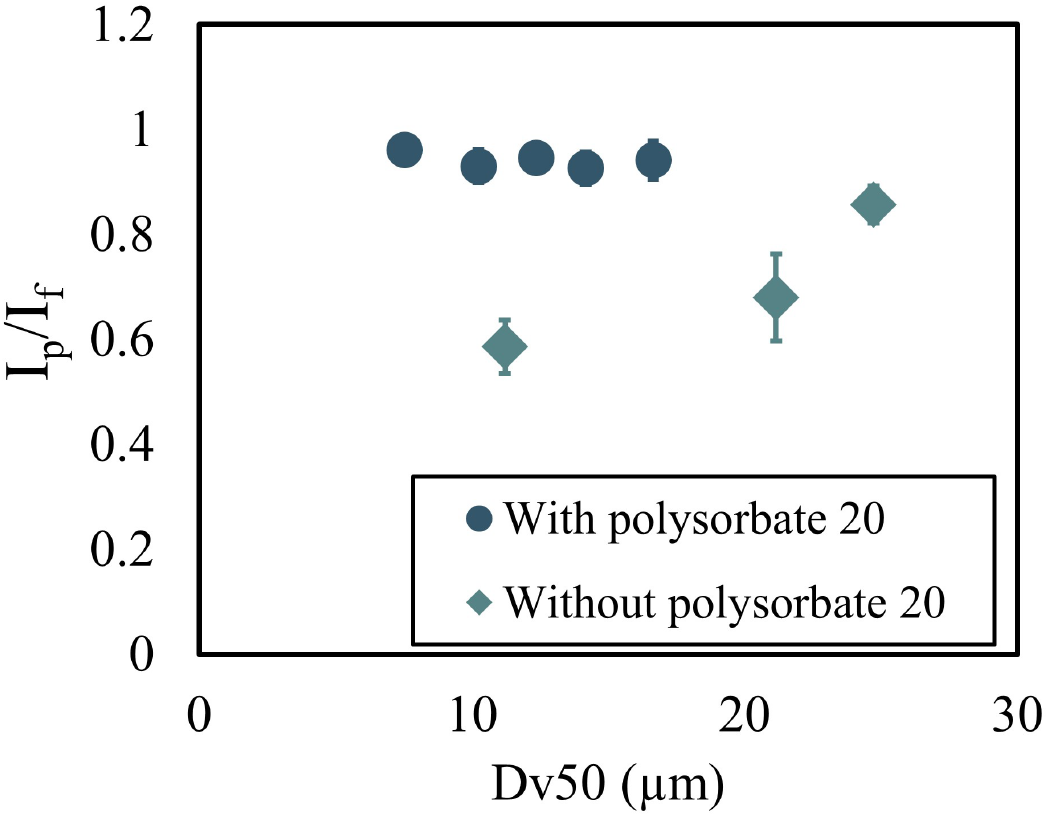
Ratio of GFP fluorescence intensity (*I*_p_/*I*_f_) in RTAD-dehydrated powders as a function of volume median droplet diameter (Dv50) of aerosol generated by RTAD atomizer, with and without surfactant polysorbate 20 in the feedstock solution. Data points represent experimental runs 6-10 (with polysorbate 20) and runs 1-3 (without polysorbate 20), as detailed in Table 1. This comparison illustrates the protective effect of surfactant on activity of GFP molecule across various droplet diameters during the RTAD process.

In contrast, with surfactant present, *I*_p_/*I*_f_ ratios remained independent of droplet diameters generated by the RTAD atomizer (Figure 8). For runs 6-10, *I*_p_/*I*_f_ remained nearly constant between 0.93-0.96, despite Dv50 more than doubling from 7.5 *µ*m to 16.6 *µ*m.

These results demonstrate GFP’s sensitivity and the protective effect of polysorbate 20 on GFP. Proteins and peptides are known to denature at liquid-gas interfaces (*27, 28*). Surfactants like polysorbate 20, common in pharmaceutical formulations, can migrate to interfaces, protecting protein molecules from interface-induced damage (*29, 30*). Also, it is known that in traditional spray drying, increasing surface-active molecules in the feedstock can eventually eliminate proteins from precipitating at spray-dried particle surfaces (*31*). Therefore, we conclude that polysorbate 20 effectively protects GFP from interfacial stress-induced damage during novel RTAD process. This protective effect is particularly evident for smaller droplet diameters, as shown in Figure 8, where the presence of surfactant significantly reduces GFP activity loss.

## 4. Summary and Conclusions

In this study we demonstrate the efficacy of using green fluorescent protein (GFP) as a sensitive model biologic for investigating thermophysical and chemical stresses induced during the dehydration of bioformulations in our novel Rapid Room-Temperature Aerosol Dehydration (RTAD) process. The fluorophore of GFP, enclosed in a beta-barrel scaffold (*6*), showed sensitivity to interfacial phenomena and recirculating flow patterns in the drying chamber, providing rapid and detailed insights into the drying process. We have shown that only a small concentration of GFP (≤ 0.03 mg/ml) is needed to generate robust fluorescence intensity measurements that can reflect the structural integrity of the protein. By analyzing the fluorescence intensity ratio of GFP after and before dehydration under different conditions, we were able to iteratively improve the RTAD system design.

Our findings reveal that interfacial and mechanical stresses during dehydration can cause irreversible changes to GFP fluorescence intensity, with variations of up to 28% observed between different chamber configurations. The presence of polysorbate 20 as a surfactant significantly mitigated the negative effects of interfacial stresses on GFP fluorescence intensity, highlighting the importance of formulation composition in preserving protein integrity during dehydration. The novel RTAD technology offers several key advantages over conventional spray drying:

1. Room-temperature operation: By leveraging sub-20 *µ*m droplets, RTAD achieves rapid evaporation at ambient temperature, significantly reducing thermal stress on sensitive biologics.
2. Enhanced preservation of biological activity: Through optimized chamber design and formulation, we achieved GFP fluorescence recovery / biological activity of 96%, indicating the preservation of protein structure and demonstrating the potential for superior product quality.
3. Rapid process optimization: GFP fluorescence measurements provide results within minutes, compared to hours required for chromatographic analysis of non-fluorescent proteins.

Our Computational Fluid Dynamics (CFD) modeling and numerical simulations elucidated the critical role of flow patterns in the drying chamber, demonstrating that longer particle residence times and flow recirculation can subject sensitive materials to additional stresses. This insight led to the development of an optimized chamber design (configuration C) that minimizes recirculation zones and provides more uniform drying conditions. The methodology presented here contributes significantly to the advancement of room-temperature dehydration technologies for thermally sensitive biologics. By using GFP as a model molecule, we were able to efficiently optimize various aspects of the RTAD process, including formulation composition, chamber design and gas-droplet-particle flow patterns. Future research directions could include investigating a broader range of biopharmaceuticals to validate the generalizability of these findings, exploring the long-term stability of dried products produced under different RTAD conditions, and developing in-line monitoring techniques based on GFP fluorescence for real-time process optimization. In conclusion, in this work we establish GFP as a powerful tool for assessing and optimizing biopharmaceutical drying processes, while demonstrating the potential of RTAD technology to revolutionize the production of dry powder biologics. The high biological activity recovery achieved (96%) underscores the promise of RTAD for preserving sensitive biological products during manufacturing, potentially opening new avenues for the development and delivery of biopharmaceuticals.

## 5. Acknowledgement

This work was supported by the National Science Foundation under STTR Phase I Grant award #2304461. The authors acknowledge the use of Princeton’s Imaging and Analysis Center, which is partially supported by the Princeton Center for Complex Materials, a National Science Foundation MRSEC program (DMR-2011750). The numerical simulations presented in this article were performed on computational resources managed and supported by Princeton Research Computing of the Office of the Dean for Research of Princeton University. We express our gratitude to Professor Robert K. Prud’homme from the Department of Chemical and Biological Engineering of Princeton University for helpful discussions and for providing access to the equipment in his laboratory. We are grateful to Dr. Mark Esposito (KayoThera), Dr. Kenric Hoegler (AstraZeneca), Dr. Rina Dukor (BioTools), Dr. Yelena Pyatski (BioTools) and Dr. Sadegh Poozesh (AstraZeneca) for their invaluable discussions and expert advice.

## Appendix A. Study of the independence of numerical solution on the mesh size

A polyhedral mesh was used for the computational domain, with boundary layer mesh cells grown from the walls to ensure a no-slip boundary condition. Adaptive mesh refinement was employed in regions downstream of the drying gas and atomizing gas inlets, where the velocity magnitude gradients were high, to capture the flow features with greater accuracy. A grid independence study was performed to ensure the accuracy of the numerical simulations. Five sets of computational grids were investigated, with total mesh cell counts of 0.70 million (M1), 1.33 million (M2), 2.51 million (M3),

3.97 million (M4), and 5.81 million (M5). We compared the change in azimuthally averaged *z*-velocity along the chamber radius at a cross section of *z* = 0.1 m for all five grids in Figure A1. As the number of mesh cells increased, the averaged *z*-velocity converged at all locations along the radius. The difference in velocity at the center between M4 and M5 was less than 2%, indicating good convergence. Therefore, the computational grid M4 was adopted for all simulations in this study.

The criteria for a converged solution were defined as residuals reaching 10^−3^ for continuity, momentum, *k*, and *ϵ*, and 10^−6^ for energy and species. Typically, convergence was achieved after 1000 numerical iterations. The computations were performed on clusters of Princeton Research Computing. A typical simulation took approximately 5 hours, utilizing 16 parallel 2.6 GHz Intel Skylake CPU cores.

**Figure A1.**
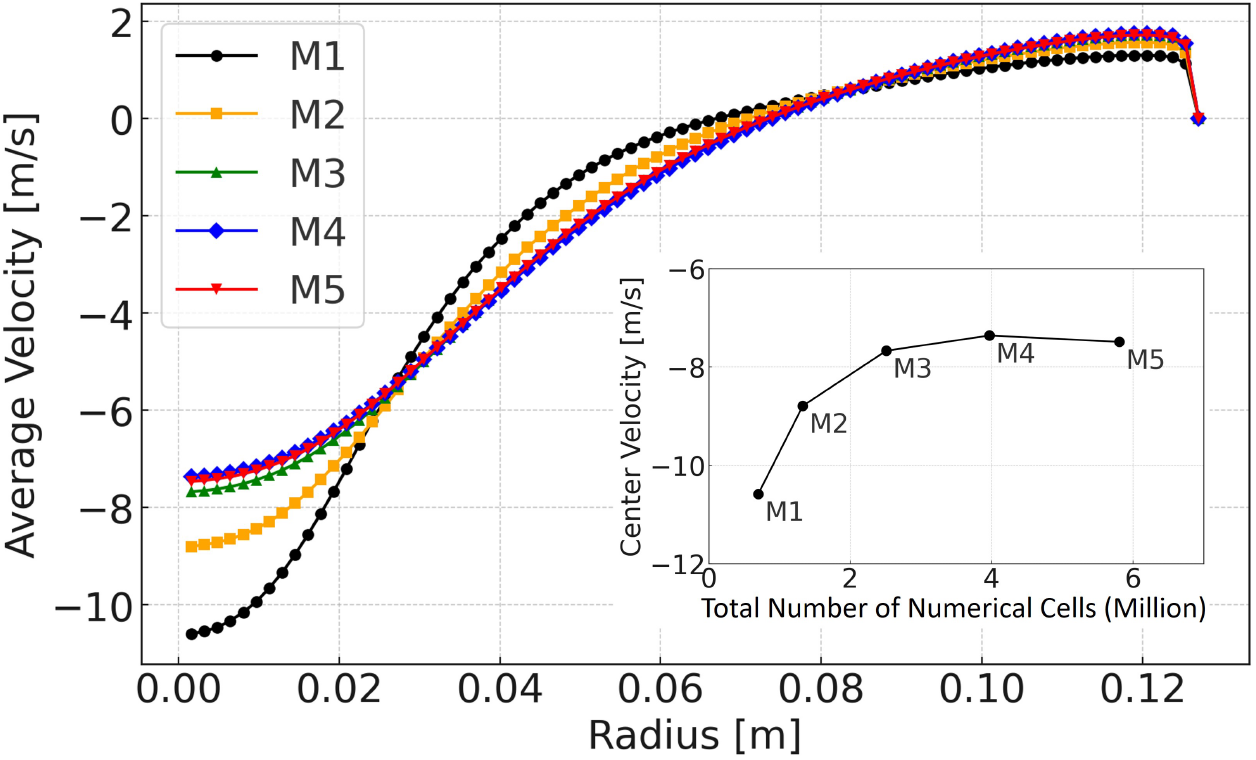
Study of numerical solution dependence on mesh size. Change in azimuthally averaged *z*-velocity along the chamber radius at a cross section of *z* = 0.1 m for numerical grids M1, M2, M3, M4, and M5. The change of *z*-velocity with the total number of cells is also presented.

